# Evolution of drug-binding residues in eukaryotic ribosomes

**DOI:** 10.1101/2024.11.28.625670

**Authors:** Lewis I. Chan, Chinenye L. Ekemezie, Karla Helena-Bueno, Charlotte R. Brown, Tom A. Williams, Sergey V. Melnikov

## Abstract

Drugs that target eukaryotic ribosomes are becoming increasingly important as research tools and potential therapies against cancer and pathogenic eukaryotes. However, in the absence of comparative studies, we currently do not know how many eukaryotes possess ribosomal drug-binding sites identical to those in humans, and how many significantly differ from humans. To address this, we traced the evolutionary history of individual ribosomal drug-binding residues from the emergence of eukaryotes to the present day. We found that ribosomal drug-binding sites are divergent across eukaryotic clades, with some of the clades exhibiting more substitutions in their ribosomal drug-binding sites compared to humans than humans do compared to bacteria. Overall, our work provides a resource for understanding the evolutionary divergence of drug-binding sites in eukaryotic ribosomes, which may inform the use of ribosome inhibitors as research tools and lineage-specific drugs against eukaryotic parasites.

## INTRODUCTION

Small molecule inhibitors of eukaryotic ribosomes are widely used as research tools and experimental medicine (1–6). With over twenty chemical families available for ribosome targeting, we can arrest ribosomes at every step of their work cycle (1). This enables the use inhibitors like puromycin, cycloheximide and others in ribosome profiling (6), and drugs like homoharringtonine and ataluren treat cancer and genetic disorders (7–9). Furthermore, ribosome inhibitors can be photocaged to halt protein synthesis with a light signal, facilitating studies of protein synthesis in neurons during memory formation (10–12).

This widespread use of ribosome inhibitors relies on the assumption that ribosomes are highly conserved, allowing commonly studied organisms, including humans, to serve as generalized model eukaryotes (1,13). However, emerging evidence indicates that ribosomes from some eukaryotes have divergent drug-binding sites and varying affinities to ribosomal inhibitors. For example, the aminoglycoside-binding site of the ribosome is divergent in the parasitic fungi Microsporidia (due to the 18S rRNA substitution A1491U) and pathogens *Leishmania* and *Trypanosoma* (bearing the substitution C1409U) (14–16). Both these substitutions increase aminoglycoside resistance (17,18). However, in the absence of comparative analyses, it is unknown whether eukaryotes with divergent drug-binding sites are exceptions or a common evolutionary trend. Consequently, we do not know how many eukaryotes have identical or different ribosomal drug-binding sites to humans.

The lack of such comparative studies originates in part from the problem of spurious mutations in biological sequences. Previous studies have shown that datasets of biological sequences may contain errors resulting from incorrect base calls during sequencing, assembly mistakes, or erroneous annotations of sequencing data (19,20). Because many biological sequences are assembled from multiple shorter reads, public repositories were estimated to contain up to 20% chimeric sequences, where different segments of a single sequence originate from distinct species rather than a single source. Other prevalent artifacts include sequencing errors, contamination from DNA of organelles, plasmids, and viruses, and the presence of pseudogenes (19,20).

This problem of a high prevalence of pseudogenes is especially acute for eukaryotes. For example, a typical bacterium like *E. coli* bears only 10-20 pseudogenes, equating to roughly one pseudogene per 300 genes (21). By contrast, the human genome bears at least 19,724 pseudogenes or roughly one pseudogene for each protein-coding gene (22). The pseudogene count is even higher in other eukaryotes, especially plants, with barley, for example, bearing 89,440 pseudogenes per 35,000 genes of its genome (23). As a result, evolutionary analyses of ribosomal genes are typically focused on larger structural changes, such as insertions or deletions, especially in bacterial species (24–29), leaving a knowledge gap about the variability of the ribosomal drug-binding residues across eukaryotes.

Here, we address this problem by implementing a novel comparative approach that resolves common problems in public sequencing data repositories—such as chimeric sequences, species misannotations and pseudogenes—thereby ameliorating the impact of database errors on our analyses and conclusions (19,20). This allowed us to compare rRNA sequences of ribosomal drug-binding residues in 8,563 eukaryotes from all major branches of eukaryotes, identify their most common variants and trace their evolutionary history throughout 2 billion years of eukaryotic history.

## RESULTS

### Mapping substitutions in ribosomal drug-binding sites during eukaryotic evolution

To help resolve the problem of pseudogenes and better distinguish signal from noise in eukaryotic rRNA datasets, we devised an approach based on similarity of rRNA sequences in closely related species on the tree of life (**Figure 1**). In this approach, we filtered out sequence variants that are present only in a single sequence, single rRNA operon, single strain, or a single species (**Figure 1a**), but not in neighboring species on the tree of life (**Figure 1b**). This allowed us to identify changes at drug-binding sites that are present in at least two closely related eukaryotic species, thereby helping distinguish false positives and very recently acquired mutations from ancient, genuine evolutionary changes that represent common features of eukaryotic clades (**Figure 1**).

**Figure 1.**
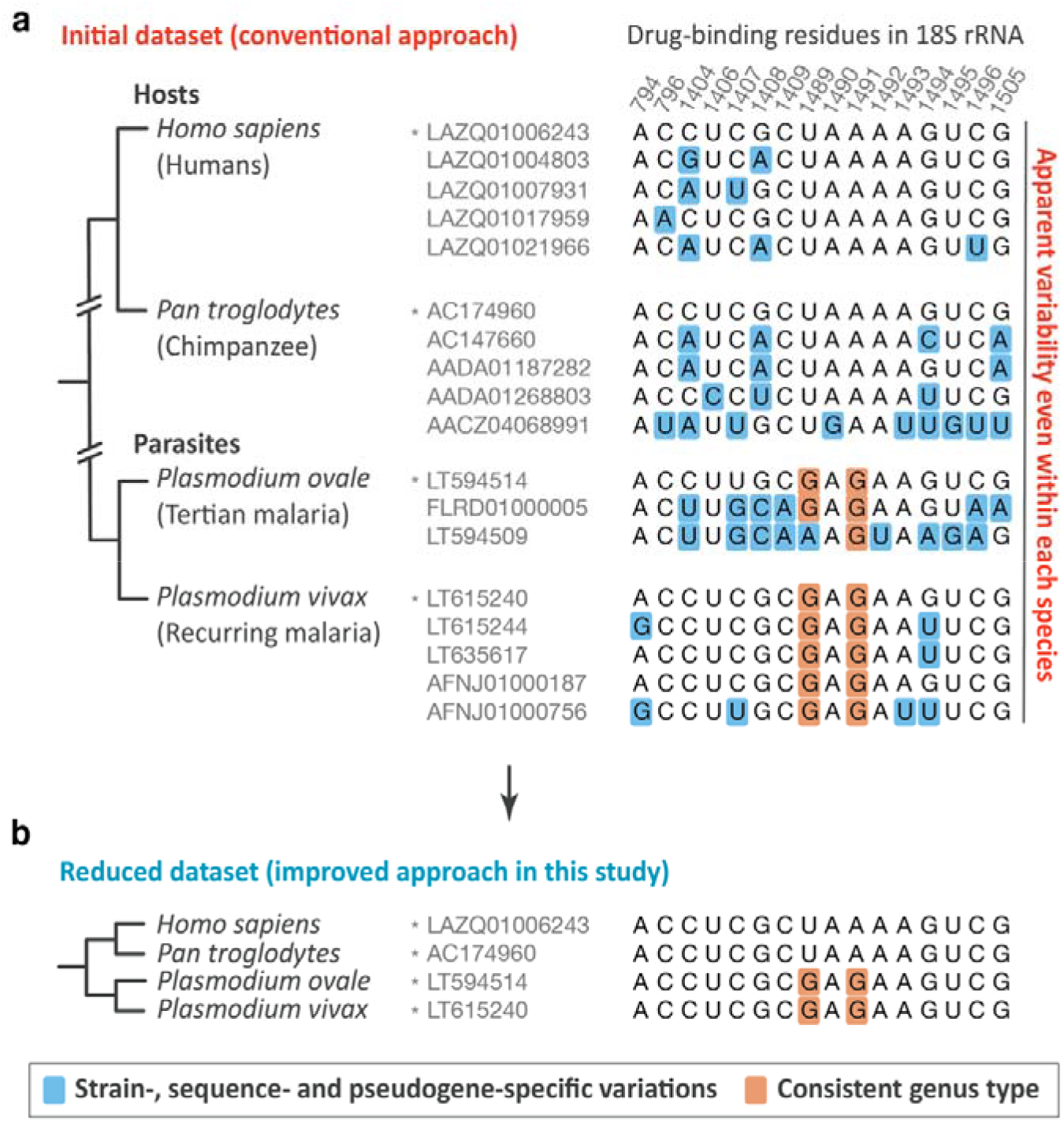
Evolutionary proximity reveals common variants at drug-binding sites in eukaryotic ribosomes. **(a)** Multiple sequence alignments compare the ribosomal drug-binding residues of the 18S rRNA from a few representative eukaryotes, including humans and chimpanzees, as well as *Plasmodium* parasites. The rRNA sequences are shown as deposited in the SILVA database, which is the most commonly used database for studying rRNA sequences across species. The panel illustrates that the ribosomal drug-binding residues appear to be highly variable even within a single species, but this seeming variability may be attributed to pseudogenes. (**b**) Multiple sequence alignment shows the reduced dataset of 18S rRNA sequences, which includes only those sequences from panel (**a**) that are conserved between immediate neighbors on the tree of life. This approach illustrates the key idea of our study: we reasoned that if a certain rRNA substitution is ancient, it will be present not only in a given species but also in its neighbor on the tree of life (e.g., other eukaryotes from the same genus). By contrast, mutations that are either most recently acquired or false positives (due to misannotations, sequencing errors, or pseudogenes) should be present only in a single sequence, single rRNA operon, single strain, or a single species, but not in neighboring species on the tree of life. This approach helps eliminate pseudogenes and reveal the most common variants of the ribosomal drug-binding sites of eukaryotic ribosomes.

Using this approach, we assessed the conservation of 58 ribosomal drug-binding residues in rRNA sequences from the SILVA database, which is the most complete database of non-redundant rRNA sequences (**Methods, Table S1**) (30). We then traced the evolutionary history and conservation of individual drug-binding residues across 8,563 representative eukaryotes from 5,148 distinct genera. Thus, we were able to reconstruct the evolution of drug-binding residues in eukaryotic ribosomes at the levels of single residue, single species and single ribosome-targeting drug (**Figure 2**,**3**).

**Figure 2.**
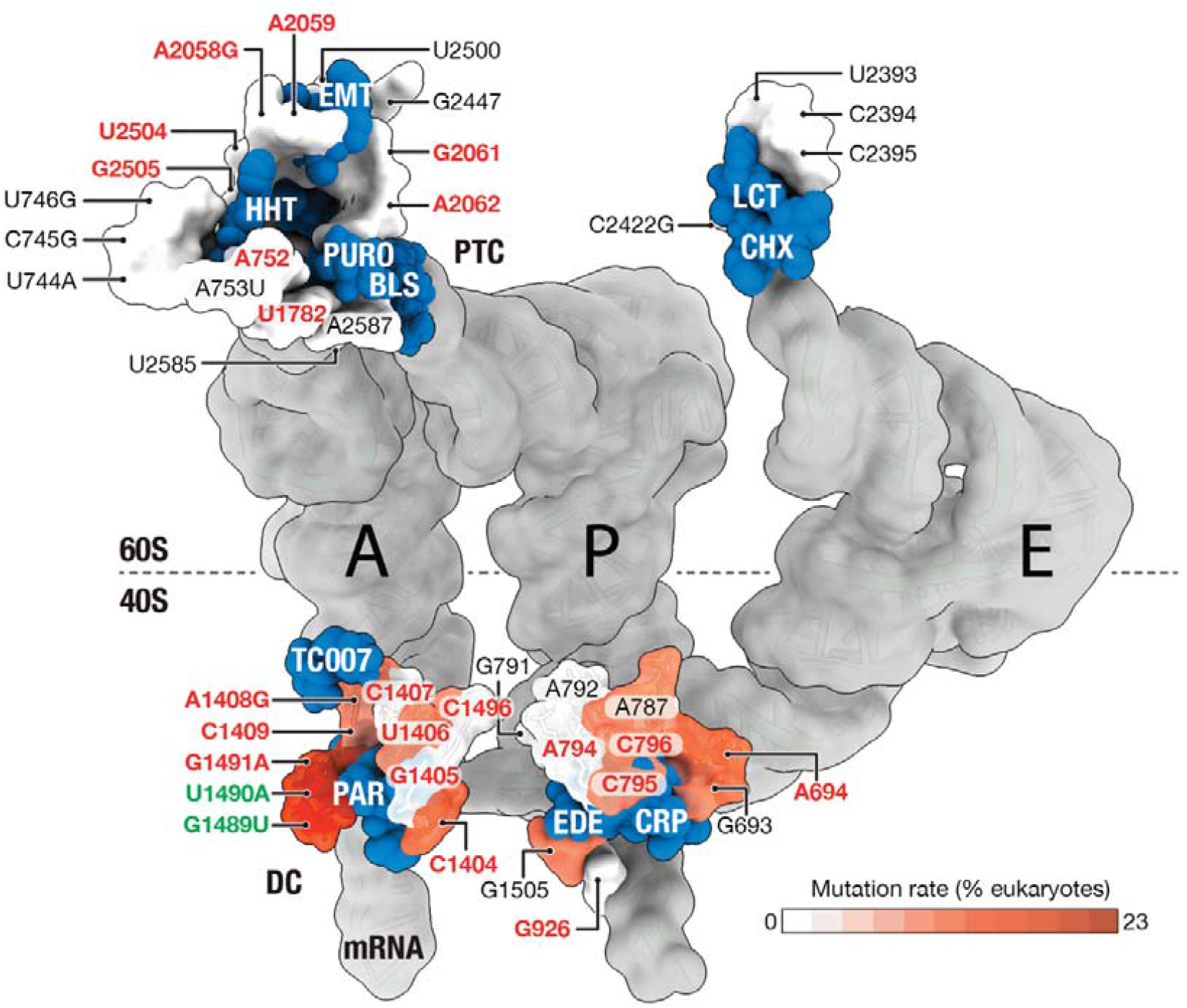
Most variable drug-binding residues of eukaryotic ribosomes. Superposed structures highlight the binding sites for ribosome-targeting drugs (in blue) in the ribosome, with each drug-binding residue of eukaryotic ribosomes colored by its conservation across the 8,563 representative eukaryotic sequences analyzed in this study. The figure represents the superposition of multiple PDB structures that capture eukaryotic ribosomes in complex with drugs, as listed in (**Table S1**). The drugs are labeled as follows: EMT – emetine, HHT – homoharringtonine, PURO – puromycin, BLS – blasticidin S, LCT – lactimidomycin, CHX – cycloheximide, PAR – paromomycin, EDE – edeine, CRP - cryptopleurine. The figure shows that, in the large ribosomal subunit, drug-binding residues are highly conserved across eukaryotes, suggesting that all eukaryotes may share a uniform recognition of drugs that target the large ribosomal subunit. However, the drug-binding residues of the small ribosomal subunit are highly variable, both in the decoding center and in the mRNA channel of the ribosome, which illustrates the frequent occurrence of species-specific features of the drug-binding pockets in the small subunit of the eukaryotic ribosome.

### Drug-binding sites of the small subunit are highly variable across eukaryotes

Our analysis showed that the large and small ribosomal subunits have strikingly dissimilar conservation of their drug-binding rRNA nucleotides (**Figure 2, SI Data 4 and 5**). The large subunit exhibits an exceptionally high degree of sequence conservation: U2393 and G2422 were the only variable drug-binding residues, having been replaced with As in the pathogenic yeast *Malassezia*, a causative agent of seborrheic dermatitis and tinea versicolor (31). Thus, the large subunit of eukaryotic ribosomes appears to have the uniform structure of the drug-binding sites.

In stark contrast, in the small subunit residues, the majority of drug-binding residues were variable, including 15 of the 26 residues in 18S rRNA. Some variations—such as A787C, C796U, and C1404U—were rare, occurring in only 20 to 40 eukaryotes within our dataset. Other variations, including U1489G/C and A1491G/C/U, were highly prevalent, each observed in nearly 2,000 species. Notably, we observed variations not only in rRNA residues that differ between bacteria and humans but in residues currently viewed as universally conserved (**Figure 2**). Thus, we found that 23% of eukaryotes harbor between one and four fixed changes in their rRNA drug-binding residues compared to humans.

### The evolution of drug-binding residues of the eukaryotic ribosome

We next traced the evolutionary history of variations for individual drug-binding residues of eukaryotic ribosomes (**Figure 3**). This allowed us to estimate the evolutionary age of these variations and their occurrence across eukaryotes. Previously, the last common ancestor of modern eukaryotes, LECA, was estimated to have emerged around 1.8-2.3 billion years ago (32,33), with the divergence of stem eukaryotes from archaea occurring somewhat earlier (2.7-2.2 Ga, (33)). Our analysis indicates that some of the key differences between bacterial and eukaryotic ribosomes in terms of drug sensitivity were already established before the divergence of eukaryotes from Archaea. For example, our analysis indicates that G1408 and A1491 in 18S rRNA (compared to A1408 and G1491 in bacterial 16S rRNA) were already established prior to the archaea-eukaryote divergence Importantly, each of these mutations has been characterized as a critical determinant for the species-specific recognition of aminoglycoside antibiotics: they prevent the binding of paromomycin in the ribosomal decoding center due to the steric clash between the rRNA base and the antibiotic (1), leading to approximately 1,000-fold lower ribosome affinity for aminoglycosides (17). Our analysis showed that the major lineages of eukaryotes accumulated additional drug-binding variants during their early diversification.

**Figure 3.**
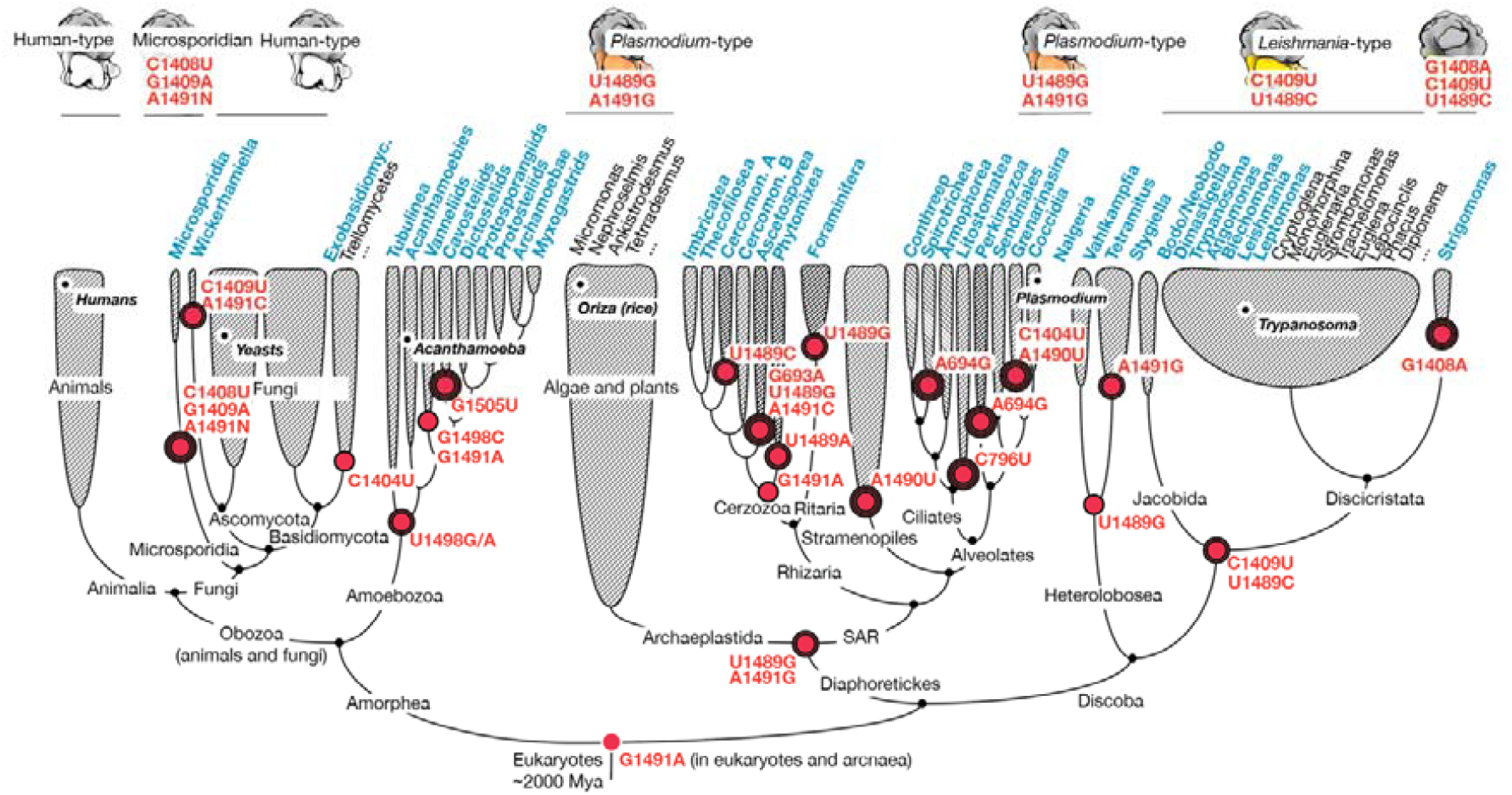
Most common variants of the drug-binding residues of eukaryotic ribosomes. The eukaryotic tree of life shows the evolutionary history of the naturally occurring variations in the drug-binding residues of the eukaryotic ribosome. Genera highlighted in blue correspond to branches including parasitic species. Red circles indicate evolutionary events related to the acquisition of substitutions in ribosomal drug-binding sites and provide an estimated age of these mutations based on the existing phylogenetic evidence listed in (**Table S3**). Overall, the figure shows that yeasts and humans, traditionally viewed as representative model eukaryotes, in fact bear relatively unusual ribosomal binding site of aminoglycoside antibiotics—due to the residue A1491 compared to G1491 in the majority of eukaryotes from non-animal and non-fungal branches. Therefore, almost all other eukaryotes have structurally distinct binding sites for aminoglycoside antibiotics, where G1491 is present instead of A1491, with some branches (e.g., Discicristata) bearing additional mutations in this and other drug-binding sites of the ribosome (e.g., C1404U, G1408A and C1409U) compared to humans.

Specifically, after Discoba separated from the other eukaryotes approximately 1.4 to 1.8 billion years ago (33), it acquired the substitution U1489C. Subsequently, most Discoba, including the branches of Jakobida and Discicristata, also acquired the substitution C1409U. As a result, these species have gained a structurally distinct drug-binding site that bears the C1409U and U1489C substitutions compared to humans. Notably, these substitutions have been previously characterized in the ribosomal structures of *Trypanosoma* and *Leishmania* species (14–16,18). Our analysis shows that these mutations are evolutionarily ancient and represent a common characteristic not only of *Trypanosoma* and *Leishmania* but of the entire eukaryotic clade of Discoba, which includes human pathogens *Naegleria fowleri* and *Balamuthia mandrillaris*.

The TSAR lineage also acquired conserved drug-binding site changes after diverging from other eukaryotes, including U1489G as well as G1491A, which reverted the sequence to make it identical to the bacterial variant, A1491. Both these substitutions have been fixed and shared by all species of this branch, including modern plants, algae, and single-celled eukaryotes from SAR species, including *Plasmodium* species and other human pathogens.

The Amorphea branch further split into the ancestor of modern fungi and animals (Obazoa), most of which have retained the ancestral ribosomal state, and the branch of amoeba-bearing Amoebozoa, which have acquired the mutation of the A1498 base shared by all members of this clade, including human pathogens *Entamoeba, Acanthamoeba* and others.

These early mutational events were later followed by the acquisition of numerous additional mutations, resulting in more than 60 unique combinations of the sequence of drug-binding residues among modern eukaryotes (**Figure 3, SI Data 5&6**).

### Certain clades of fungi bear altered ribosomal drug-binding sites compared to humans

Our further analysis of the eukaryotic tree of life showed that, although most animals and fungi share conserved drug-binding residues, certain fungal lineages have gradually acquired multiple changes at these sites compared to humans (**Figure 4**). This was observed, for example, in the deep-branching fungal lineage Microsporidia, all members of which are fungal parasites of animals, including humans (34). In Microsporidia, the initial mutation likely occurred shortly after the split into *Nematocida* species, which retained human-type (that is, ancestral) drug-binding sites, and the remaining microsporidia, which acquired a mutation in the decoding center at residue A1491, which has been identified as a key determinant of species specificity for aminoglycoside antibiotics (17,1). Subsequent evolution of microsporidia has led to additional mutations in the decoding center, including C1489U in the *Vavraia* branch and G1048A in species of *Enterocytozoon*. Hence, most microsporidia exhibit a more dissimilar decoding center compared to humans than humans compared to *E. coli*.

**Figure 4.**
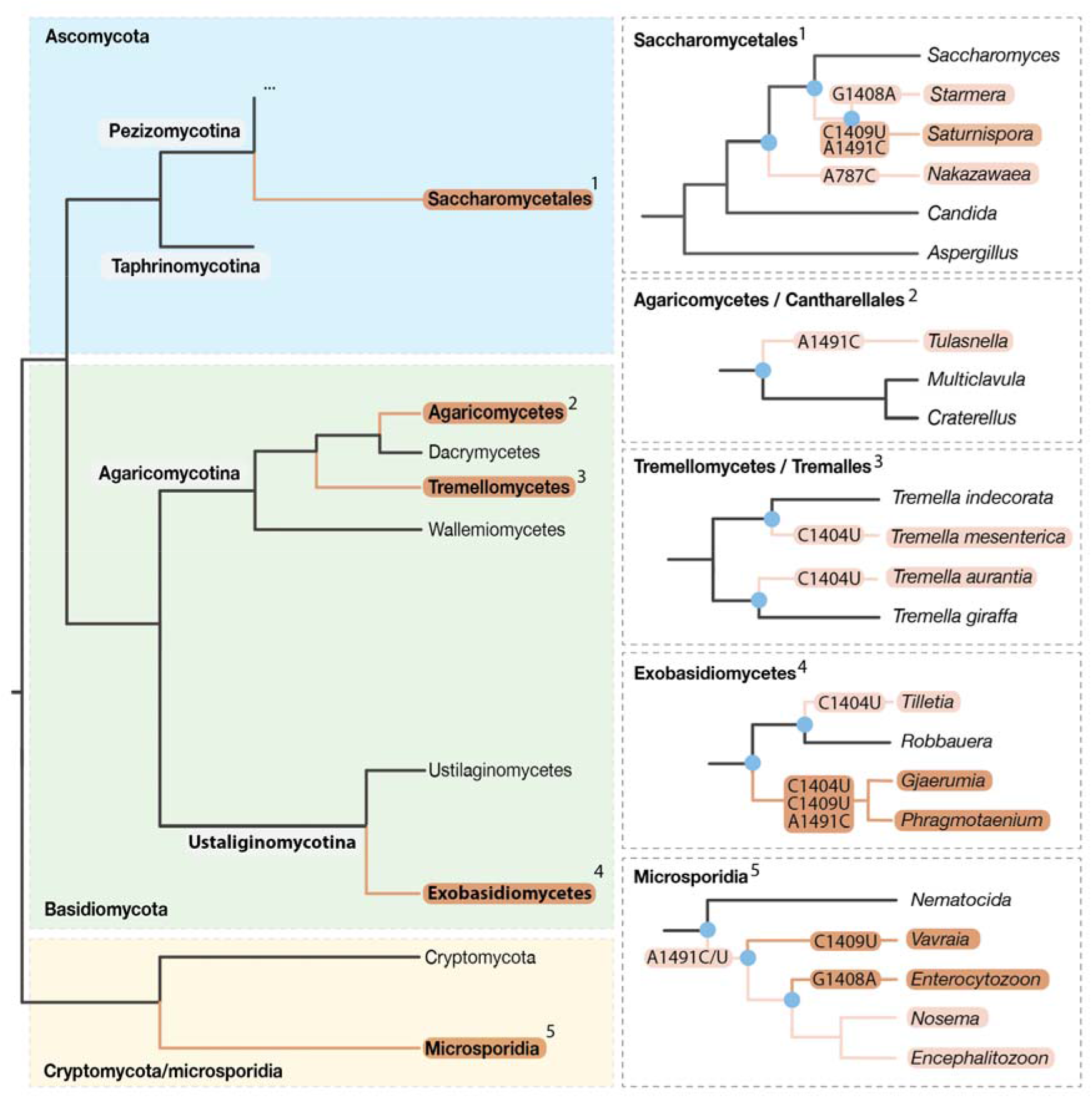
Many fungi bear derived ribosomal drug-binding sites compared to humans. The schematic structure of the fungal tree of life shows that, after separating from animals, some lineages of fungi, such as Microsporidia and Saccharomycetales, continued to diversify their ribosomal drug-binding residues. In total, of the 1,253 genera of fungi analyzed in this study, 101 (∼7%) were found to carry one or more mutations in their ribosomal drug-binding residues compared to humans.

The remaining fungal lineages were further split into several clades, including Ascomycota and Basidiomycota, with specific branches within these clades acquiring mutations in both the aminoglycoside-binding site and the mRNA channel. These branches include the fungi *Trellomycetes* and *Exobasidomycetes* (C1409U), which encompass pathogenic organisms, such as *Tilletia horrida* (C1404U), a common rice pathogen, and *Gjaerumia minor* (C1404U, C1407U, A1494C), implicated in keratitis of humans. Overall, among the 1,201 fungal species analyzed in our study, we found approximately 7% to exhibit dissimilar ribosomal drug-binding sites compared to humans, due to variations in the 18S rRNA bases A787, C1408, C1409 and A1491 (*E. coli* numbering is used throughout the manuscript. Please refer to Table S2 for the correspondence between species) within the decoding center and the mRNA channel of eukaryotic ribosomes.

Thus, we found that most fungi have identical ribosomal drug-binding residues compared to those in humans, suggesting that ribosome-targeting drugs are unlikely to serve as broad-spectrum antifungal agents. However, species from certain fungal clades, particularly microsporidia, possess multiple mutations in their drug-binding residues compared to humans, indicating the potential for their safe lineage-specific targeting.

## DISCUSSION

### Animals and fungi possess distinct ribosomal drug-binding sites compared to most other eukaryotes

In this study, we have assessed the evolutionary history and conservation of ribosomal drug-binding residues across the eukaryotic domain of life. We determined the natural variations for each of the 58 ribosomal drug-binding residues in 8,563 representative eukaryotes, enabling us to identify lineages with dissimilar residues compared to humans. One major finding of our work is that yeasts and humans, traditionally used as model organisms for studying ribosome-targeting drugs in eukaryotes, exhibit rather unusual variants of drug-binding residues compared to most other eukaryotes. Specifically, yeasts and humans, along with most Amorphea species, appear to have kept the ancestral sequence variant of the drug-binding sites, whereas the other branches of eukaryotes have acquired substitutions in the drug-binding residues G1408, C1409, A1491 of the 18S rRNA. Importantly, some of these substitutions have reverted the rRNA sequences of eukaryotic rRNA to its bacterial-type variants (e.g. in the 18S rRNA residue G1491 in Diaphoretickes).

One important implication of this finding is how we currently study eukaryotic ribosome targeting with small molecules. From the perspective of drug sensitivity, ribosomes from different organisms are often separated into two major groups: bacterial-type and eukaryotic-type (13). This separation is based on the overall protein content and the presence of rRNA expansion segments, allowing the use of organisms like *E. coli* or *T. thermophilus* as representative bacteria and organisms like yeasts or humans as representative eukaryotes (1). However, our work shows that this division, while helpful in many studies of eukaryote-specific ribosomal proteins and rRNA expansion segments, is incomplete when applied to the drug-binding residues of the eukaryotic ribosome. For instance, some groups of eukaryotes, such as SAR, share more similarities in their drug-binding residues with bacteria than with humans due to the convergent evolution of the 18S rRNA variation A1491G. Additionally, some others, like microsporidia, are highly dissimilar to both bacteria and humans. Instead, our work shows that the simplistic division of ribosomes into bacterial and eukaryotic is mostly accurate only when applied to the large ribosomal subunit. By contrast, a range of different residues, and so potentially drug-binding sites and sensitivities, are found across eukaryotes in the small subunit of the ribosome, including (roughly) an animal/fungi-type, *Leishmania*-type, and *Plasmodium*-type, among many others.

### Many of the naturally occurring variations in ribosomal drug-binding sites predate the origin of antibiotic-producing bacteria

Since when have eukaryotic ribosomes started to diversify their drug-binding residues? And what were the evolutionary forces driving this diversification of ribosomal drug-binding residues? Our mapping of the rRNA sequence variants on the tree of life allows us to gain insights into both of these questions. According to this analysis, substitutions at the drug-binding sites of eukaryotic ribosomes began to accumulate from the origin of the first eukaryotic branches >1.3 Ga (33), with some mutations mapping to the stem lineages of major groups including Amorphea, Discoba and TSAR. This means that common 18S rRNA variants such as A1491G in SAR, and G1409A and C1489U in Discoba species, emerged significantly earlier than the antibiotic-producing genus *Streptomyces*, which is responsible for most natural ribosome-targeting drugs known to date and is estimated to be 382 million years old (35). This analysis suggests that the evolution of drug-binding residues in nature, at least during early eukaryotic evolution, differs from their evolution in clinical settings and has likely been driven by factors other than the existence of antibiotics in the environment.

### Implications for species-specific targeting of eukaryotic ribosomes

Previous studies showed that mutations in individual rRNA drug-binding residues can confer drug resistance by reducing drug-affinity as much as 1,000 fold (36). In this study, we showed that some of these variations, previously observed in clinical isolates of human pathogens or laboratory-engineered resistant strains, are widely present in nature.

Our analysis, along with the previously obtained experimental studies, suggests that some of the naturally occurring variations are likely neutral. Specifically, variations of residues G1489U and U1490A have been characterized as neutral for ribosome targeting by aminoglycoside drugs, as demonstrated through ribosome mutagenesis in *Mycobacterium smegmatis* and *E. coli* (37–39).

However, other variations, including 13 residues of the 18S rRNA, have been shown to alter the shape of the drug-binding pocket and define species-specific ribosome targeting with drugs (red labels in **Figure 2** and **Table S4**). For example, the variations A1408G and G1491A in the ribosomal decoding site have been characterized as determinants of species-specific binding of aminoglycoside antibiotics, allowing for the safe targeting of bacterial ribosomes (A1408 and G1491) without affecting ribosomes in humans (G1408 and A1491) (40). Experiments in yeasts showed that when eukaryotic ribosomes are mutated to introduce the bacteria-specific variant G1491 in the 18S rRNA (instead of A1491), they develop aminoglycoside sensitivity similar to *E. coli*, evidenced by a 60-fold higher sensitivity to the antibiotic paromomycin. Similarly, mutating the G1408 site to the bacteria-type A1408 resulted in a ∼200-fold increase in sensitivity of yeast ribosomes to the antibiotics neomycin and kanamycin A (17). Our analysis reveals that these variants are not limited to bacteria, but are also prevalent across eukaryotes, naturally occurring in up to 23% of eukaryotic species.

Overall, our findings present both concerns and opportunities for using drugs to target eukaryotic ribosomes. We show that the drug-binding sites of eukaryotic ribosomes often exhibit significant variability across different eukaryotic lineages, thus necessitating caution and additional analyses prior to applying ribosome-targeting drugs to non-model eukaryotes. On the opportunity front, our work suggests the possibility for lineage-specific ribosome-targeting drugs across a wide variety of eukaryotes. Notably, we demonstrate that structural variants identified in *Leishmania* and *Plasmodium*, previously considered idiosyncratic to rare eukaryotic lineages, are in fact common characteristics of entire eukaryotic supergroups. What’s more, some lineages of eukaryotes have more dissimilar ribosomal drug-binding sites compared to humans than humans do compared to bacteria. Since eukaryotic pathogens pose significant and emerging global health challenges, our work suggests that novel therapeutics can be found among new generations of ribosome inhibitors that target lineage-specific substitutions in the rRNA.

## METHODS

### Locating the drug-binding residues in eukaryotic ribosomes

To identify the ribosomal residues that are directly involved in drug recognition, we analyzed 29 previously determined structures deposited in the Protein Data Bank (41) in which eukaryotic ribosomes were bound to a representative of each family of ribosome-targeting drugs (**Table S1**). Using ChimeraX, we selected non-hydrogen atoms of the ribosome located within 4.3 Å from a drug. These sets of atoms were then reduced to only include atoms that belonged to amino acid side chains of ribosomal proteins and aromatic bases of rRNA. This allowed us to determine the coordinates of each ribosomal residue that makes a sequence-dependent contact with ribosome-targeting drugs, as well as the numbering correspondence between conserved rRNA residues of *E. coli, S. cerevisiae* and *H. sapience* ribosomes (**Table S2**).

### Creating a high-fidelity dataset of rRNA sequences for evolutionary analyses

To enable accurate analysis of rRNA sequences, we used the SILVA database and have devised a multi-step data reduction strategy in order to bypass the notorious issues of automated data annotation and sequencing, including the occurrence of chimeric sequences, misannotated phylogeny, non-genomic DNA and pseudogenes (**SI Figure 1**). In brief, as our starting point we used the SILVA’s dataset NR99 v138.1, which contains 510,508 16S/18S rRNA sequences corresponding to 78,104 unique “species” (including metagenomic variants, unknown species, species candidates and other ambiguously defined species) (“SSU dataset”) and 95,286 sequences of 23S/5.8S rRNA from 20,471 unique “species” (“LSU dataset”) from all domains of life (30). We first reduced these SILVA datasets by eliminating sequences from non-eukaryotic species as well as from unidentified eukaryotes for which we could not confirm the identity of an organism. This included sequences whose headers contained the words “unidentified”, “uncultured”, “metagenome”, “cluster” and “sp.”, as well as sequences whose names began with a lowercase letter. The obtained datasets were then further reduced by eliminating rRNA sequences that are encoded by plasmids and organelles. We then removed all sequences of apparently poor sequencing quality, based on the presence of ‘ambiguous’ sites represented by symbols such as “R”, “X”, “N”, and “S” instead of the standard set of RNA bases “A”, “G”, “C”, and “U”. The sequences from each of these datasets were initially aligned using MAFFT v7.490 with default settings (FFT-NS-2) (42) during the quality control analysis. The sequences that passed the quality control were then re-extracted from the original SILVA dataset of aligned rRNA sequences and are shown in (**SI Data 2, 3**).

### Assessing the conservation of the ribosomal drug-binding residues

To simplify the assessment of conservation of drug-binding residues of the eukaryotic ribosome, we trimmed the aligned rRNA sequences to only the drug-binding residues. This file was then manually inspected for quality control, and the rRNA with apparent truncations in the drug-binding sites (based on the presence of gaps in the place of drug-binding residues) were removed from our further analysis. The remaining sequences were then reordered according to the location of species on the tree of life. The resulting dataset was further reduced to a single sequence per organism using the following rule: whenever the same organism contained several sequence variants, we used only one variant, which was identical (or the most similar) to the sequences of the adjacent members on the tree of life (e.g. other eukaryotic species from the same genus). This allowed us to identify mutations that were a common characteristic of a clade of species and eliminate sequencing and annotation artifacts, as well as mutations that were characteristic of an individual strain or an individual rRNA operon (**SI Data 4, 5**).

### Reconstructing the evolutionary history of mutations in ribosomal drug-binding sites

To trace the evolutionary history of sequence variations in the ribosomal drug-binding residues, we used 16S rRNA sequences listed in (**SI Data 5**) to generate a phylogenetic tree. The tree was generated using default settings on Mega11 (43). The tree was then colored by conservation of ribosome-drug-binding residues by highlighting species and lineages with the human-type and non-human-type sequences of ribosomal drug-binding residues (**Figures 3**,**4**). The relative age of each mutation was estimated using TimeTree (44) or the original studies cited where appropriate in the manuscript.

## ACKNOWLEDGEMENTS

We would like to thank Robert Hirt, Claudia Schneider, Wyatt Yue and Heath Murray (all Newcastle University, UK) and Erik C. Böttger (University of Zurich, Switzerland) for their comments on the manuscript and Yuri Polikanov (University of Illinois at Chicago, USA) for encouraging this study. This work was supported by the BBSRC Doctoral Training Program (BB/T008695/1 to L.I.C. and C.R.B.), the MRC Discovery Medicine North Doctoral Training Partnership (MR/N013840/1 to C.L.E.) and the Newcastle University Academic Track Fellowship (to S.M.).

## Notes

https://doi.org/10.6084/m9.figshare.27918987

### Competing Interest Statement

The authors have declared no competing interest.

### Summary of Updates

We improved the clarity and brevity of this work.

